# Manifold properties in the macaque medial premotor cortex during switching from attending to tapping to a metronome

**DOI:** 10.1101/2025.10.07.681062

**Authors:** Dobromir Dotov, Abraham Betancourt, Jorge Gámez, Hugo Merchant

**Affiliations:** Department of Biomechanics, University of Nebraska Omaha, Omaha, NE; Instituto de Neurobiología, UNAM, Campus Juriquilla, México; Now at California Institute of Technology, Pasadena, CA

## Abstract

Animals synchronize their movements with external rhythms to coordinate perception and action, but the neural population mechanisms that allow them to attend and then initiate and sustain these rhythms remain unclear. Using high-density recordings from the medial premotor cortex (MPC) of two macaque monkeys, we investigated neural dynamics during an attend-then-synchronize tapping task with visual metronomes. We found low-dimensional neural manifolds that capture neural population trajectories. During the attention phase, trajectories exhibited increasing amplitude and oscillatory modulation along successive stimuli, consistent with a resonant-like mechanism. Transition to tapping was marked by a reliable shift into a distinct manifold subspace, enabling accurate decoding of the switch in behavior. In addition, large amplitude and oscillatory indices were higher in successful tapping synchronization trials than in incorrect ones. These findings demonstrate that macaque MPC activity evolves along smooth, low-dimensional trajectories whose geometry governs different perceptual and motor aspects of tapping synchronization.

## Introduction

Animals coordinate their actions with predictable events in their surrounding environment to perceive, move, detect danger, and communicate with each other (Merchant et al., 2009; Merchant, Crowe, et al., 2011; Romo et al., 1996). For communication, different forms of rhythm exist in different species as a function of their vocalization capacities and ecological demands (Greenfield, 1983; Legett et al., 2021; Ravignani et al., 2014; Snedden et al., 1998). Group synchronization can confer a stability advantage to the rhythmic performance of individuals (Dotov et al., 2022). Notably, **beat-based rhythm** in music and dance affords synchronous interactions between participants (Merchant et al., 2015a) and engages sensorimotor synchronization that is linked with complex loops of attention, perception, and action (Patel & Iversen, 2014; Pérez et al., 2023; Repp, 2005). **Beat perception** allows predicting or anticipating future events (Harding et al., 2025; Merchant et al., 2014) by co-opting the motor system (Cannon & Patel, 2021; Merchant & Honing, 2014; Morillon & Baillet, 2017; Zatorre et al., 2007). The human audiomotor system is designed to extract the temporal patterns from continuous streams of sounds and to perceive a steady pulse or beat in music spontaneously (Garcia-Saldivar et al., 2024). The beat is a periodic internal representation that is projected into the future to predict when the next event is likely to occur, and to trigger the initiation of movements so that they coincide or anticipate this cognitive event (Rajendran et al., 2025). Crucially, there is evidence that an internal or covert beat is present in a variety of rhythmic tasks (Lenc et al., 2021; Repp et al., 2008; Repp & Su, 2013; Toiviainen et al., 2010). In perceptual tasks, it can be detected in large-scale brain activity as EEG oscillations of prominent amplitudes at the frequencies of the beat (Nozaradan, 2014; Nozaradan et al., 2018). This is the case for both weakly and strongly periodic auditory stimuli (Lenc et al., 2021; Nozaradan et al., 2012). How beat-based rhythm can harness populations of neurons is still debated and several explanatory mechanisms have been proposed. In this work, we first briefly review these mechanisms, and then we use high-density neural recordings from macaques to investigate how population dynamics emerge while attending to and tapping along an isochronous stimulus.

### Neural entrainment and neural resonance

One of the influential models in this context, neural entrainment, studies how an external rhythm can phase-synchronize intrinsic brain oscillations and focus sensory processing of stimuli that occurs on the beat (Lakatos et al., 2008; Schroeder et al., 2008). This shares certain features with dynamic attending theory, according to which the synchronization of neural oscillations to external stimuli, such as speech and music, creates a rhythm of attention with enhanced and suppressed processing at salient points (Large & Jones, 1999). Entrainment models can be characterized specifically by the implication that intrinsic self-sustained oscillations must preexist the stimulus so that they can be entrained by it, see Figure 1A (Schroeder & Lakatos, 2009). Neural resonance is an alternative mechanism in which amplitude and oscillation dynamics are not intrinsic but emerge as a result of the rhythmic input of stimulus energy, see Figure 1B (Başar et al., 1992; Doelling & Assaneo, 2021; Helfrich et al., 2019; Large & Kolen, 1994). Mathematical models show that populations of weakly coupled excitable units, driven by resonance phenomena, can exhibit time-keeping properties (Izhikevich, 2007; Zemlianova et al., 2022).

**Figure 1.**
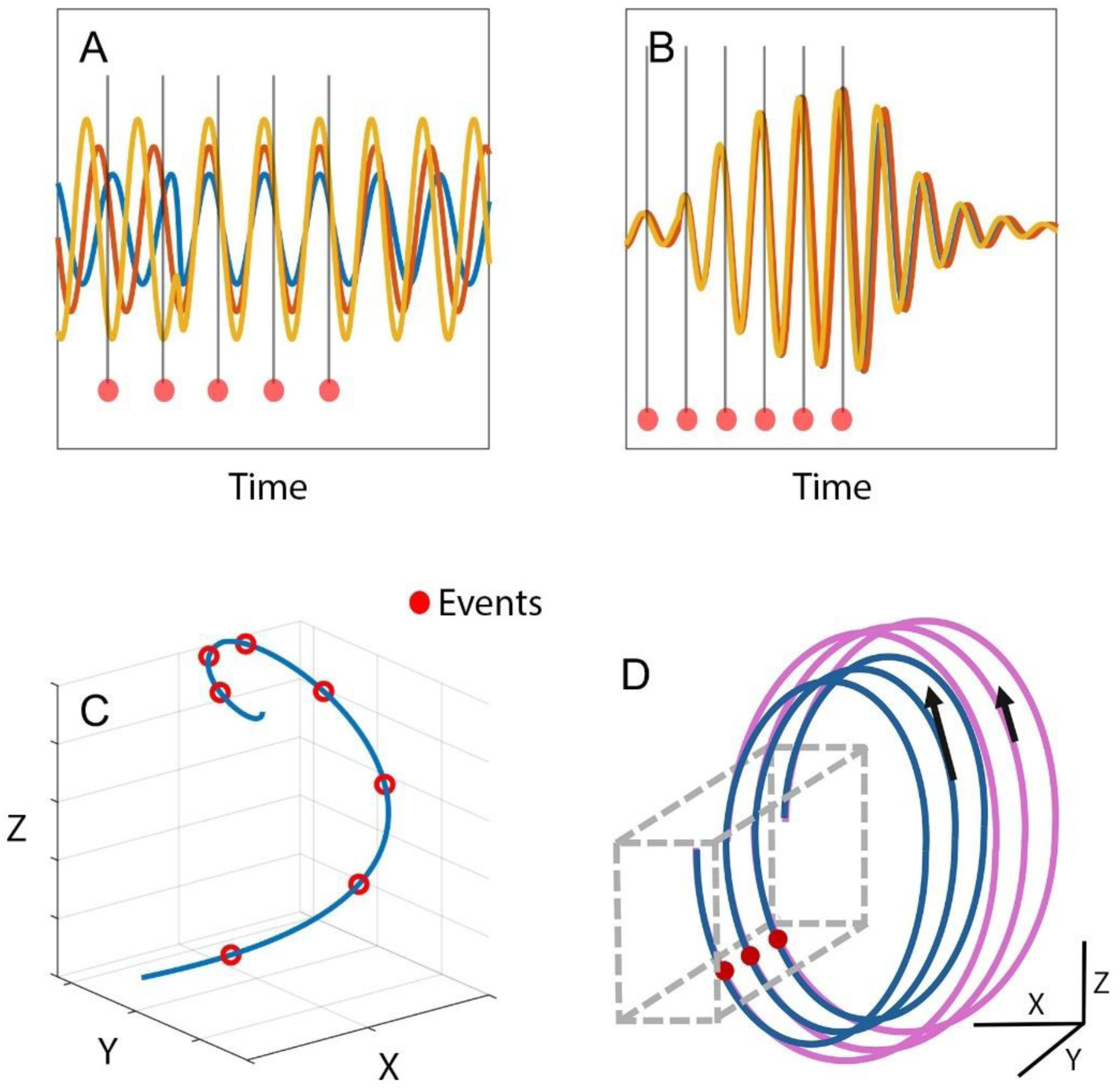
Schematic illustrations of different but not mutually exclusive mechanisms for organizing neural populations during synchronization with rhythmic events. For the sake of generality, events marked as red dots could be either stimuli or motor events. A) Entrainment is characterized by alignment of intrinsic on-going oscillations phase-shifted by periodic events (S1-S5), represented by the vertical lines. The intrinsic oscillations change phase until they are phased-locked by S5. B) Resonance is characterized by emerging oscillations with increasing amplitude when the events have a frequency close to a preferred frequency of the neural dynamics. The intrinsic oscillations increase in amplitude and reach a maximum by S5. C) A low-dimensional neural state trajectory unfolds along the duration of the trial. Presumably, the dimensions X, Y, and Z are on a task-related manifold that is embedded inside the high-dimensional space of neuronal firing rates. D) Similar to C), low-dimensional neural state trajectories during a rhythmic tapping task with either of two different intervals, shown in blue (short) and pink (long). The amplitude and speed (tangent vectors) are candidate parameters for encoding the interval duration (Betancourt et al., 2023; Gámez et al., 2018). Tap times are found within a convergent section of the state space, shown as a dashed box.

### Neurophysiology of synchronization to a metronome in the Rhesus monkey

Questions about the nature of neural dynamics in rhythmic tasks can be addressed in detail by studying Rhesus monkeys. Their rhythmic abilities and phylogenetical closeness to the human audio-motor system make them an interesting model species to trace the emergence of rhythmic skills (Honing & Merchant, 2014). For the past 20 years, our lab has been showing that macaques have all the necessary audiomotor machinery to perceive and synchronize to the beat (Merchant et al., 2024). EEG studies in the Rhesus monkey have shown that macaques produce evoked potentials linked to the detection of isochronous auditory patterns (Honing et al., 2018), as well as to subjectively accented 1:2 and 1:3 rhythms from auditory metronomes (Ayala et al., 2017; Criscuolo et al., 2023). In addition, monkeys trained on tapping tasks can flexibly and predictively produce periodic intervals in synchrony with auditory and visual metronomes (Betancourt et al., 2023; Gámez et al., 2018), and can continue tapping isochronically without sensory cues (Bartolo et al., 2014; Zarco et al., 2009). Notably, they tap consistently to the subjective beat of music excerpts with different tempos, choosing freely a tapping phase for each song (Rajendran et al., 2025). Furthermore, monkeys show an error correction mechanism during synchronization that helps them control the duration of the produced intervals, that is more efficient at the preferred tempo of the animals (Castillo-Almazán et al., 2025). Therefore, these observations suggest that during tapping synchronization, monkeys use a complex rhythmic timing mechanism that includes both attentional and predictive components.

Imaging studies in humans have shown that the neural substrate of beat-based timing includes pathways within the voluntary motor control system, including medial premotor areas (MPC: SMA and pre-SMA), basal ganglia (most often the putamen, but also caudate nucleus and globus pallidus), and the motor thalamus, all of which form one of the cortico-basal ganglia-thalamo-cortical loops (Grahn & Rowe, 2009; Kung et al., 2013; Sánchez-Moncada et al., 2024). In the macaque, recent neurophysiological studies indicate that the MPC is involved in the internal pulse while tapping to a metronome (Merchant, Zarco, et al., 2011; Merchant & Averbeck, 2017). Cells in an MPC population are recruited in rapid succession, producing a progressive pattern of activation. This flexibly fills the beat duration and indicates relative interval progressions (Crowe et al., 2014; Merchant et al., 2015b).

### Neural population trajectories

The advent of high-density neural recordings using implanted electrodes in animal models has enabled a closer examination of the relationship between individual neurons and population dynamics in a given area of the cortex (Berényi et al., 2014; Buzsáki et al., 2012; Mendoza et al., 2016a). Populations of neurons combine sparse coding, where few neurons are strongly tuned to a given stimulus feature, and mixed selectivity, where neurons are tuned to multiple statistically independent behavioral parameters. As a result, the same population can respond flexibly to different stimuli (Panzeri et al., 2015). Of interest to us, this suggests that we need to study how the same neural population responds to different intervals and different phases of the task.

Central to the present work is the substantial interest in the fact that the many neurons recorded in a high-density setup can embed a low-dimensional state space when the individual cells are correlated. This is uncovered with dimensionality-reduction techniques that take as input the high-dimensional set of individual activities and extract a low-dimensional set by projecting them on fewer variables and leaving out redundancies (Cunningham & Yu, 2014; Shenoy et al., 2013). The reduced variables define a neural state space, often characterized as a smooth manifold inside the original high-dimensional space (Jazayeri & Ostojic, 2021). Neural manifolds contain the possible states of a population of neurons, given the combination of intrinsic constraints, such as anatomy and neurophysiology, and extrinsic constraints, in the form of behavioral dynamics (Perich et al., 2025). The geometry and kinematics of the state trajectory through the manifold can encode behavioral events, see Figure 1C, and its parameters correspond to parameters such as interval duration (Egger et al., 2019; Sohn et al., 2019), reaching for a target (Laje & Buonomano, 2013; Stine & Jazayeri, 2025; Zhou et al., 2022), and switching between waiting and execution (Kaufman et al., 2014). In premotor and motor areas, population dynamics compute preparation parameters and trigger targeted movements (Churchland et al., 2010; Pastor-Bernier et al., 2012).

### Population trajectories in rhythmic tasks

It is known that neural trajectories often exhibit a rotational component (Churchland et al., 2012; Kaufman et al., 2014). Yet, most studies of state space manifolds have been done in the context of single-shot behaviors, such as waiting and reaching, leaving fewer opportunities to study manifolds in rhythmic tasks. In simulations, it is possible to train recurrent neural networks to exhibit oscillations on a low-dimensional reduced space (Mastrogiuseppe & Ostojic, 2018; Zemlianova et al., 2024). In the macaque’s rhythmic tapping and synchronization with a stimulus, cyclic evolution is evident when the time-varying activity of MPC neurons is projected onto a low-dimensional state space. The population trajectories exhibit a list of interesting properties, summarized schematically in Figure 1D (Gámez et al., 2019). They have recurrent or regenerating loop dynamics for each produced interval; they converge to a narrow state space at the time of tapping, resetting the beat-based clock at this point. This internal representation of rhythm could be transmitted as a phasic top-down predictive signal to sensory areas; and geometric features such as speed and amplitude encode the produced interval duration. Furthermore, we showed that this neural population chronometer is bimodal, having the same dynamic properties in response to both auditory and visual metronomes (Betancourt et al., 2023).

This study investigates how said neural trajectories arise when a rhythmic stimulus is present, even before tapping begins. To this end, we employed the same overall animal training and testing framework and data stream, but we focused on the neural trajectory before the onset of tapping. Specifically, we employed a task with two epochs: a perceptual epoch in which the monkeys attended to the metronome stimulus without moving, and a tapping epoch in which the monkeys synchronized their movements with the stimulus. The neural population trajectories in MPC showed two different state space manifolds, one for each task epoch. In addition, there was a gradual increase in their amplitude, and the robustness of their oscillation of each loop was linked to the sequentially produced intervals.

## Methods

### Subjects

Two monkeys (*Macaca mulatta*, M1, male, 10 kg, 10 yoa, M2, female, 7 kg, 8 yoa) performed the experiments after extensive training. All the animal care, housing, and experimental procedures were approved by Ethics in Research Committee of the Universidad Nacional Autónoma de México and conformed to the principles outlined in the Guide for Care and Use of Laboratory Animals (NIH, publication number 85-23, revised 1985). The two monkeys were monitored daily by the researchers and the animal care staff to check their conditions of health and welfare.

### Task

Unlike one-shot actions, such as reaching and single-interval timing, rhythmic tapping presents the unique opportunity to measure parameters of the oscillatory dynamics of the neural trajectory. The monkeys were trained to perform an attend-then-synchronize-task (AST) (Gámez et al., 2018), which is used to investigate neural population dynamics during rhythmic tasks (Betancourt et al., 2023; Gámez et al., 2019). The animals were trained to attend to and then tap in synchrony with a sequence of discrete stimuli, see Figure 2. Animals were presented with periodic visual events. Each trial consisted of a sequence of between nine and eleven events with a fixed inter-stimulus interval (ISI). The animal had to withhold moving for at least three intervals before they could begin tapping in response to the stimuli. Trials were terminated if the animal started tapping too early, too late, or with a large asynchrony. ISI duration was a trial condition, with six conditions from 450 to 950 ms. M1 performed 1941 trials, 38% correct. M2 performed 2151 trials, 72% correct.

**Figure 2.**
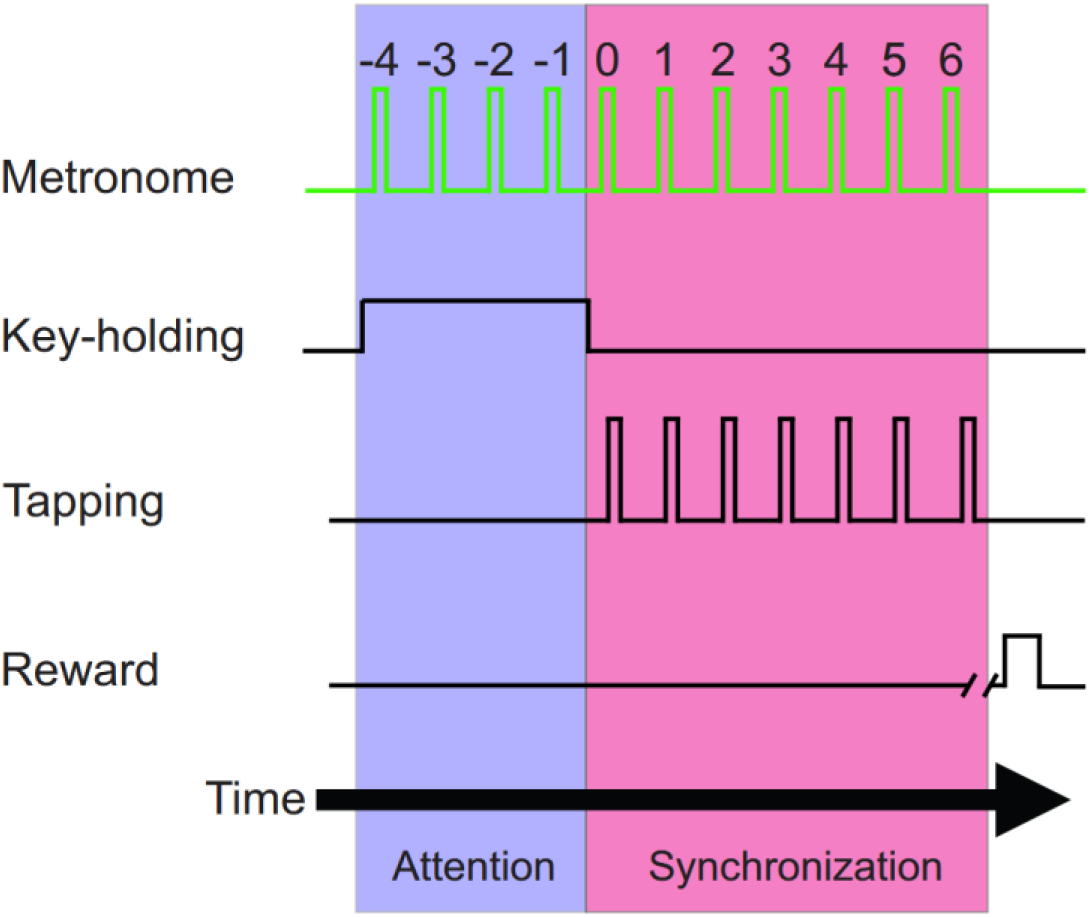
Schematic illustration of the task, which consisted of two stages. In the first stage, the animal watched the stimulus (attention) while attending to the three intervals of the metronome. The animal maintained her responding hand on a key. Then, in the second stage, the animal started tapping a button in synchrony with the metronome for six intervals to obtain a reward (synchronization).

### Neural recording

We used high-density electrode arrays developed at our lab (Mendoza et al., 2016b) and equipped with the 64-site Buzsaki64-Z64 probe. Two arrays were inserted semi-chronically in medial areas of the premotor cortex (MPC), targeting the supplementary motor (SMA) and pre-supplementary motor areas (pre-SMA), one in the left hemisphere and one in the right hemisphere.

### Spike processing and firing rates

Unit activity was extracted from the continuous neural recordings using spike sorting algorithms. For M1 we used our custom sorting pipeline (Rossant et al., 2016). For M2 we used KiloSort 2 (Pachitariu et al., 2016).

A timeseries of neuronal firing rates was computed in each trial by convolving the spike series with a Gaussian kernel (σ=10 ms, sampling rate of 1000 Hz). Trials from matching conditions on the same recording day were averaged. There were 402 cells in M1 and 1744 cells in M2 after rejecting units with low discharge rate and poor signal, the criteria being average firing rate lower than 2 spikes and cell not sampled in all conditions.

### Neural state space trajectories

The spike rate time courses of all neurons sampled in a recording day were projected to a trajectory on a low-dimensional manifold. The raw population state at a point in the trial was an *N*-dimensional vector comprising the activities of *N* cells at that moment. Separately for each monkey, we obtained a low-dimensional state-space by using principal component analysis to compute the dimensions of orthogonal variability within the embedding space and retaining only the subspace of top components (Cunningham & Yu, 2014). The projection matrix was global, meaning that it was computed after pooling all neurons from all days. Data from each trial was re-sampled and time-normalized by the ISI. This means that time was expressed in stimulus cycles, and all trials contained 50 samples per stimulus cycle, and time was shifted to set to zero the stimulus before the onset of tapping. This projection pooled statistical power from the full population of units recorded across trials to infer the underlying attentional and stimulus synchronizing state on individual trials. Furthermore, this trial-wise approach promoted the analysis of the neural trajectory along the multiple stimulus and tapping cycles of the trial.

### Analysis

We parameterized the manifold along the length of the trial by using a trial-continuous or cycle-by-cycle approach. Parameters of the population trajectory were computed using the stimulus onset and the beginning of tapping times as reference points. We analyzed how features of this trajectory evolved along the length of the trial over successive stimulus intervals because we were interested in the role of attending to the stimulus and tapping along repeated stimuli. We analyzed separately the amplitude, phase, and degree to which the neural trajectory obeyed an oscillatory dynamic. We used linear effects models separately for each monkey and each variable to test for effects of stimulus cycle number, stimulus interval duration, and correct performance.

*Amplitude* dynamics was defined as the distance of the trajectory at a point in time from the global mean. The rate of change of the amplitude was measured by taking its slope with respect to time, separately in the attention and tapping phases of the trial.

*Phase* dynamics was measured cycle-by-cycle by finding peaks of activity in individual dimensions using a peak-picking algorithm and determining their phase relative to the closest stimulus.

*Index of oscillation*. Preliminary observations suggested that each dimension of the population trajectory exhibited an asymmetric relaxation-like nonlinear oscillation characterized by slow smooth dynamics at the peak or valley and fast cusp-like dynamics at the opposite end. We accounted for the geometry of these oscillations, without modeling their generative dynamics, by fitting a function with the appropriate features. We defined an index of oscillation as the goodness-of-fit metric 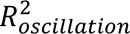 of a model with smooth oscillations and sharp peaks, (see Figure 3). The model, *ŷ* = (*a*_0_ + *a*_1_*t*) + (*b*_0_ + *b*_1_*t*)(1 − |*cos* (*πft* + *πf*/2 − *θ*/*f*)|), had free parameters for period, time-varying baseline, time-varying amplitude of the oscillation, and phase of its peaks relative to *t*=0, the stimulus after which tapping started. It was fitted using nonlinear least squares optimization separately in the attention and tapping trial sections and in each principal component. After fitting the model, the 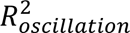 was computed independently within each successive stimulus interval of the trial to get a time-series of the index of oscillation for each stimulus interval along the length of the trial.

**Figure 3.**
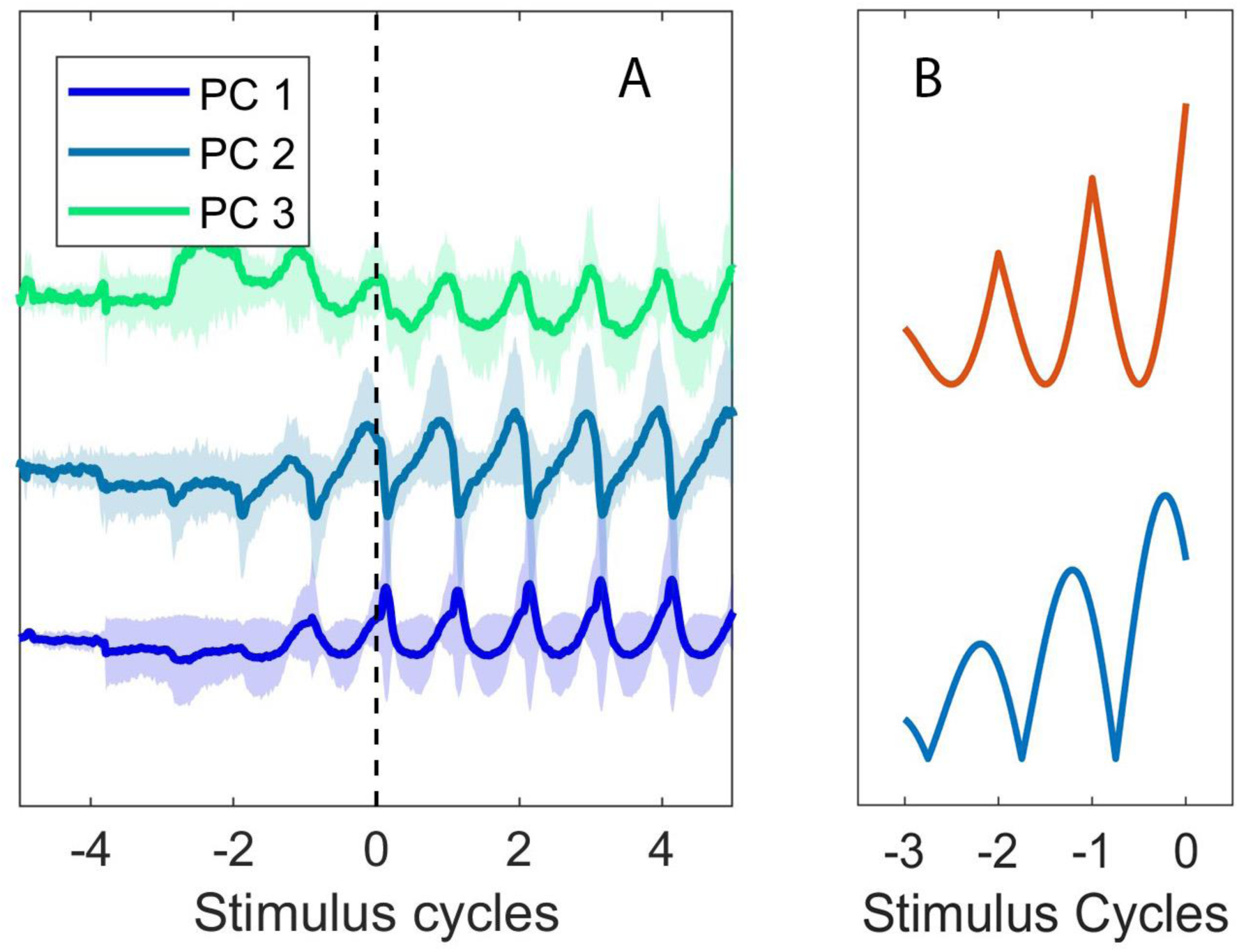
Representative population dynamics from Monkey 2 (correct trials, ISI=950 ms). A) The first three individual dimensions of the neural trajectory dynamics (Mean±SD). Time is normalized to unit stimulus. Stimulus=0 marks the onset of tapping. B) The fully embedded neural trajectory, color-coded for time, along with trial events. B) Examples of two possible functions of time realized by the oscillation amplitude model, which was used to fit individual dimensions of the neural trajectory dynamics.

*Dimensionality*. It is possible that the neural state space spans smooth high-dimensional geometry instead of a low-dimensional manifold. The former corresponds to a single scaling function that fits the full variance spectrum of the principal components (Stringer et al., 2019). We tested the hypothesis that a small number of top components, some of them with oscillatory dynamics, had higher eigenvalues than expected, indicating they diverged from the high-dimensional geometry of background activity. We combined two techniques iteratively: regressing out a variance scaling power law from the principal component eigenvalues and clustering the residuals with *k*-means to identify and remove outliers. This was inspired by a method for separating the periodic and aperiodic contributions to the EEG power spectrum (Donoghue et al., 2020).

We also considered nonlinear dimensionality reduction as suggested by recent literature (Durstewitz et al., 2023; Mitchell-Heggs et al., 2023; Stringer & Pachitariu, 2024). We used nonlinear principal component analysis with a multi-layer perceptron which, like PCA, can project unseen data to the learned low-dimensional space (Scholz et al., 2008). We did not observe qualitative differences in subsequent analyses. The interpretability of latent manifolds computed with nonlinear techniques can be difficult, yet their size is an informative parameter. We computed the intrinsic dimensionality of the manifold with a nonlinear method to serve as a lower boundary reference for the embedding dimensionality found with the linear method (Jazayeri & Ostojic, 2021).

*Behavioral state decoding*. We evaluated whether the behavioral states of attending and tapping could be classified reliably from the neural state. We trained a decoder on the top principal components selected with the dimensionality analysis. We used a support vector machine with a non-linear polynomial kernel, optimized using cross-validation, and tested on 20% of the reserved data. The decoder can be seen as a plane that separates the manifold into two regions, see Figure 4B. It classifies the behavioral state at a given point in time based on which side of the plane the trajectory is found at that point.

**Figure 4.**
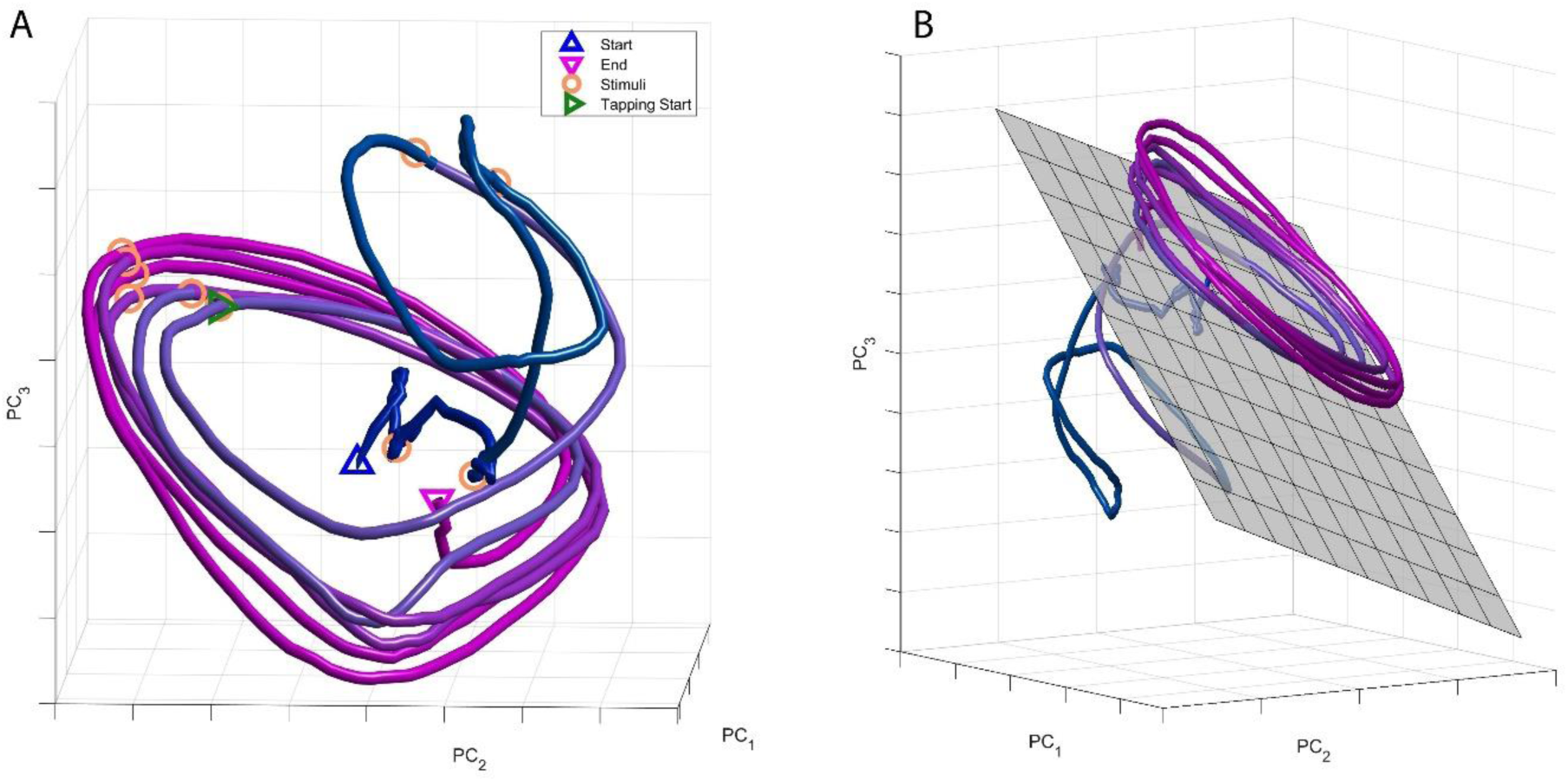
A) Representative population dynamics from Monkey 2 (correct trials, ISI=950 ms). B) The same dynamics shown from a different perspective along with the classification plane marking the transition from attention to tapping synchronization.

## Results

### Dimensionality

The size of the principal component subspace embedding the neural trajectory manifold was *d*=6 in M1 and *d*=3 in M2. These dimensions were retained for subsequent analysis. Using a nonlinear approach, we found that the correlation dimension was *d=*4.01 for M1 and *d*=3.05 for M2. As expected, these are about the same or lower than the ones obtained from the PCA variance spectrum.

### Amplitude

The time course of the neural trajectory demonstrated a pattern of rising amplitude oscillations during the attending phase, see Figure 3 for individual dimensions and Figure 4 for the fully embedded trajectory. During the attending phase of the trial, the population trajectory moved onto a spiral-like trajectory with an increasing amplitude over successive stimulus intervals, see Figure 4. The rate of change of amplitude in that part of the trial was statistically higher than zero [β=.006, *t*=3.71, *p*<.001 for M1 and β=.013, *t*=6.27, *p*<.0001 for M2], Figures 5A and 6A. In both monkeys, the effects of ISI and all interactions were not significant [*p*’s n.s.]. In the tapping phase, the rate of change was negative [M1: β=-.017, *t*=-3.93, *p*<.001, M2: β=-.010, *t*=-2.57, *p*<.05].

**Figure 5.**
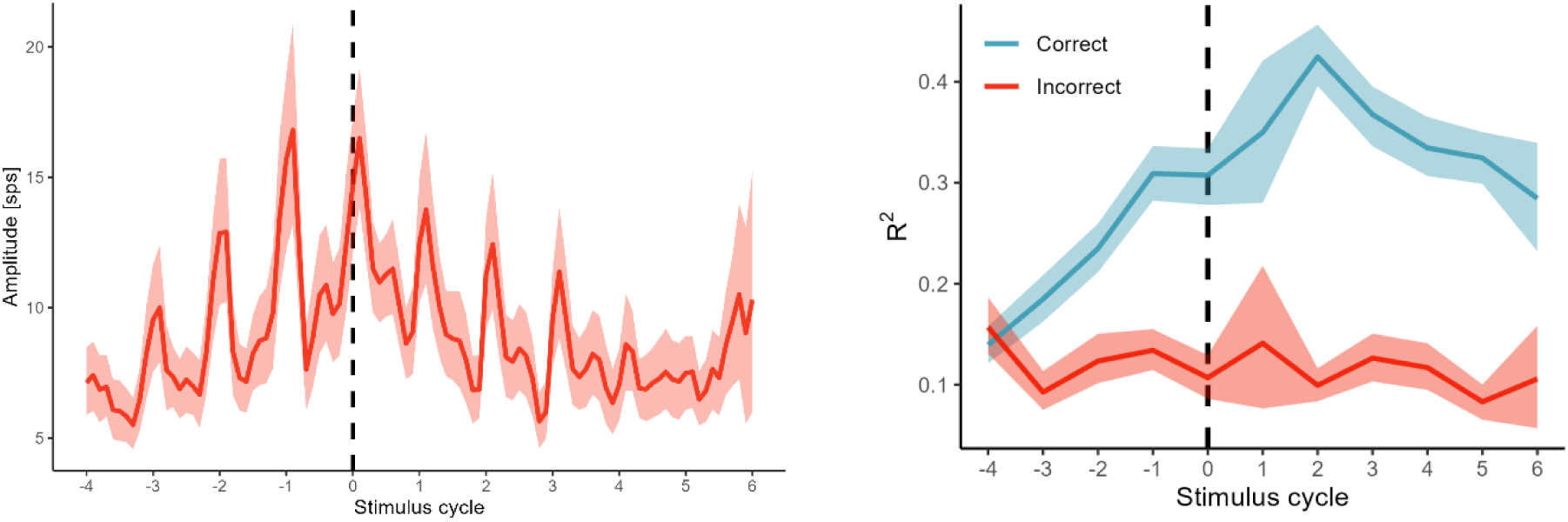
Parameters describing the population dynamics in Monkey 1. A) Amplitude of the neural trajectory, averaged across recording sessions (Mean±SD, correct trials). B) The index of oscillation averaged across manifold dimensions and sessions (Mean±SD), shown separately for correct and incorrect trials.

**Figure 6.**
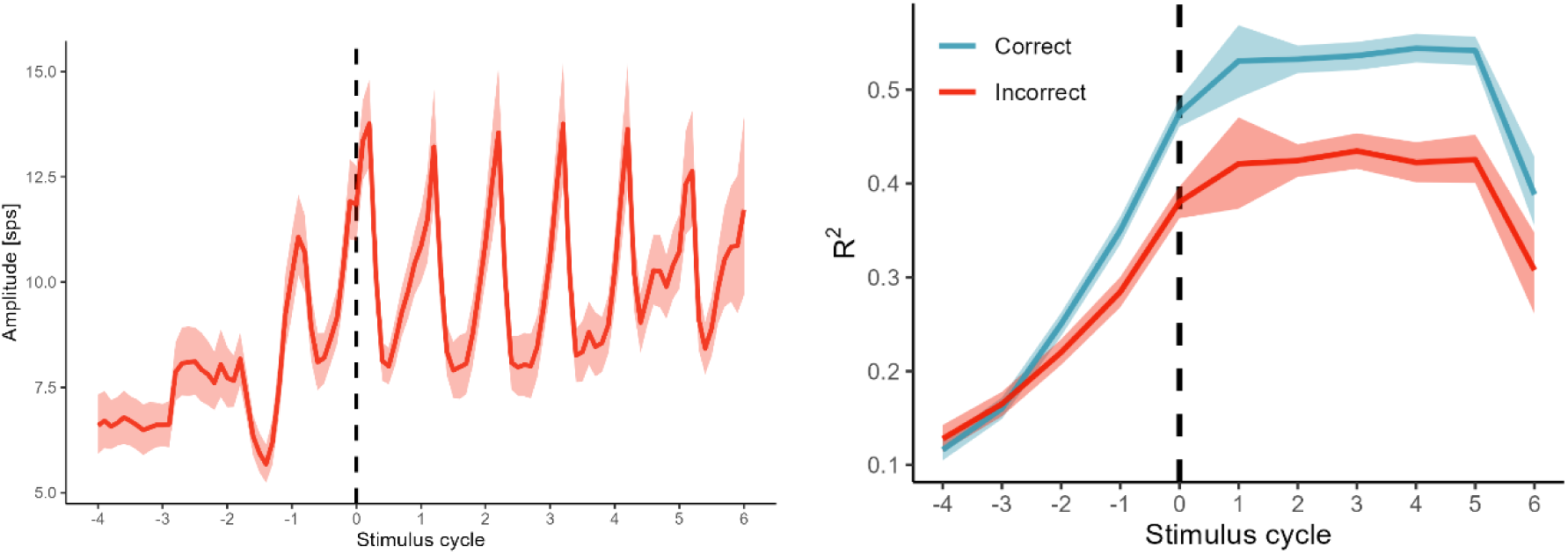
Parameters describing the population dynamics in Monkey 2. A) Amplitude of the neural trajectory, averaged across principal components and recording sessions (Mean±SD, correct trials). B) The index of oscillation averaged across manifold dimensions and sessions (Mean±SD), shown separately for correct and incorrect trials.

### Index of oscillation

Another form of evidence for the emergence of oscillatory dynamics was obtained by the time-resolved index of oscillation defined as the interval-by-interval goodness-of-fit of the oscillation function shown in Figure 3B. As can be seen in Figures 5B and 6B, the index increased during the attending part of the trial, peaked around the time of tapping onset, and it was lower in incorrect trials. Indeed, the slope of the index with respect to successive stimulus cycles in the attending part was significantly greater than zero [M1: β=.023, *t*=14.64, *p*<.001, M2: β=.074, *t*=55.65, *p*<.001]. The index of oscillation was lower in incorrect trials [M1: β=-.094, *t*=-4.34, *p*<.001, M2: β=-.046, *t*=-4.42, *p*<.001] and it was lower in higher principal components [M1: β=-.011, *t*=-6.04, *p*<.001, M2: β=-.014, *t*=-11.83, *p*<.001]. In the tapping part of the trial, the slope with respect to successive stimulus cycles was negative [M1: β=-.025, *t*=-6.69, *p*<.001, M2: β=-.009, *t*=-4.55, *p*<.001]. The index of oscillation was lower in incorrect trials [M1: β=-.262, *t*=-8.86, *p*<.001, M2: β=-.118, *t*=-6.54, *p*<.001] and in higher principal components [M1: β=-.021, *t*=-8.28, *p*<.001, M2: β=-.042, *t*=-31.05, *p*<.001].

### Phase

The phase of peaks of individual components relative to the stimuli showed little dependence on trial conditions. In the attending part of the trial, phase tended to increase with successive stimuli [M1: β=.017, *t*=4.18, *p*<.001, M2: β=.054, *t*=16.31, *p*<.001] and there were no significant associations with the other conditions [all *p*’s n.s.]. In the tapping part of the trial, phase also tended to increase [M1: β=.030, *t*=4.46, *p*<.001, M2: β=.026, *t*=4.96, *p*<.001]. It was higher in incorrect trials in M2 [β=.047, *t*=5.73, *p*<.001]. This was not a significant effect in M1 [β=.013, *t*=1.20, *p*=.23].

### Decoding

As Figure 4B shows, the population trajectory moved into a different region as the trial shifted from attending to tapping. This is consistent with the idea that neural state spaces have different subspaces for different aspects of a task. To confirm this, the classifier was able to distinguish between the attending and tapping parts of the trial solely based on the selected top dimensions. The accuracy/precision/recall were 89%/92%/88% for M1 and 92%/90%/96% for M2. This part of the analysis was restricted to data from correct trials.

## Discussion

In this study, we recorded high-density MPC neural activities during a rhythmic synchronization task performed by macaques. We found that the population dynamic was consistent with a low-dimensional neural state trajectory on a manifold with interesting properties. In the attention part of the trial, the trajectory had an increasing amplitude and increasingly coherent oscillations, presumably driven by the periodic stimuli. Concordantly, in the macaque, increasing modulation with successive stimulus intervals has been reported in populations of cells in rhythm-related deep structures such as nuclei of the basal ganglia and cerebellum (Kameda et al., 2019, 2023). In the transition from attending to tapping, the trajectory reliably switched into a different region with a stable amplitude. Dynamic parameters, particularly the index of oscillation, were positively associated with correct trial performance during this epoch. Altogether, these findings contribute to existing evidence that the geometry of low-dimensional manifolds serves to implement task-related computations such as timing and sequencing (Buonomano & Maass, 2009; Carnevale et al., 2015; Chung & Abbott, 2021; Laje & Buonomano, 2013; Schwartz & Moran, 1999; Vyas et al., 2020).

Beyond their role in timing, the geometry of neural manifolds is linked to various behavioral and cognitive processes. For example, in the prefrontal cortex of non-human primates, decisions are represented by attractor basins, where the depth and steepness of these basins correlate with decision consistency and confidence (Wang et al., 2023). Within the same region, a low-dimensional neural embedding tracks the physical properties of an object, serving as a neural substrate for mental simulation during a physical reasoning task (Rajalingham et al., 2025). In the hypothalamus of mice, a neural line attractor encodes the intensity of aggressive states related to specific actions, such as sniffing and dominance mounting (Nair et al., 2023; Vinograd et al., 2024). Recent studies also demonstrate that neural manifolds can evolve within distinct planes corresponding to different task properties. For example, in the motor cortex, neural trajectories exhibit rotational dynamics to produce reach movements that unfold within different planes corresponding to different task conditions (Sabatini & Kaufman, 2024). Similarly, in the human hippocampus, learning to perform an inference task produced the emergence of an abstract and disentangled representation of task variables on a neural manifold, with the context of the task encoded orthogonally to the stimulus (Courellis et al., 2024). All these results highlight the importance of the geometry of neural manifolds in a multitude of behaviors and brain regions.

How do these results inform the dynamic mechanisms involved in initiating and sustaining coordination with a rhythmic stimulus? We identified three non-mutually exclusive perspectives on the generation of rhythm-tracking population dynamics. It is debatable whether large-scale oscillations arising in response to a rhythmic stimulus result from the entrainment of endogenous oscillations or from a stimulus-evoked transition between different dynamic regimes (Haegens, 2020). The gradual increase in amplitude and index of oscillation reported here is consistent with resonance, more so than entrainment, yet we cannot rule out that either of these is involved at some level in the generation of the neural trajectory.

We take advantage of these considerations to re-focus the discussion on questions pertaining to the comparative abilities of humans and other animals. As reviewed above, macaques are capable of complex rhythmic perception and production skills. Yet, the animals perform such tasks for short trials, rarely exceeding ten consecutive stimulus/tap cycles. Is this related to the dynamic nature of the neural trajectories reported here? The weaving, spiral-like shape is a nonlinear property, and nonlinear dynamic systems are inherently poised to produce self-sustained oscillations. The possibility of such oscillations is an intriguing but potentially paradoxical idea. A self-sustaining oscillator in MPC state space may mean that an animal has trouble exiting that subspace to stop the behavior and initiate a new behavior. In humans, in some circumstances self-sustained oscillations of brain dynamics can be maladaptive dynamic regimes associated with pathological states in humans such as epileptic seizures (Jiruska et al., 2013), Parkinsonian tremor and freezing (Hammond et al., 2007), and schizophrenia (Uhlhaas & Singer, 2010). The trajectories reconstructed in the macaque MPC appeared to be non-tangled, meaning that they do not recur in a strict sense, although testing this was not our objective here. This would suggest that MPC neural trajectories enabling the macaque’s rhythmic tapping are not oscillators in a classical sense. Instead, the entire trial was driving the neural trajectory along a non-repeating, non-tangled path through state space where the sequence of cycles encoded the order of stimuli.

Another way of asking whether the population dynamics was self-sustaining is by considering whether it was exclusively motor and stimulus driven (Merchant and de Lafuente, 2024). Here, it did not appear to be motor driven because it started to increase during the attention phase of the trial. Yet, it also returned toward the initial state as soon as tapping stopped, see Figure 4. Hence, ambiguity remains about whether the observed dynamics in the macaque MPC can sustain a task-related rhythm that can unfold indefinitely long without ongoing stimulation. Nevertheless, in a previous study we found that the circular neural population trajectories can be maintained for three internally driven continuation intervals during the classical synchronization continuation task (Gámez et al., 2019; Merchant, Zarco, et al., 2011).

### Bursting activity is involved in coordinating individual neurons

In the future, theoretical models need to address how individual neurons affect or define the dynamics in low-dimensional manifolds linked to a task. In doing so, models need to reconcile the fact that the fine-scale neurophysiology of brain oscillations is distinct from the smooth dynamics observed at the population scale. Cortical network oscillations constitute synchronized changes in excitability levels of neuronal circuits, with frequencies spanning from 0.01 to more than 200 Hz (Buzsáki & Draguhn, 2004). The local field potentials (LFP’s) are voltage recordings which reflect the cumulative electrical activity of populations of neurons within 250 microns of the tip of an extracellular recording electrode. The oscillatory power of LFP’s directly impacts the phase and probability of an individual neuron’s discharge time and the discharges of functionally related neurons are grouped during the excitatory phase of distinct oscillations. Neural assemblies grouped by oscillations can discharge to downstream targets which require multiple active presynaptic synchronic inputs to produce an action potential (Abeles, 1982).

Oscillations provide windows of opportunity in which neurons are sensitive to inputs via fluctuations between hyperpolarized and depolarized phases of neuronal membrane potentials. The phase of waves of activity can be coordinated with external stimuli (Buzsáki & Draguhn, 2004) or with other internal neural processes to achieve large-scale integration of information (Kayser et al., 2009; Singer, 2018; Varela et al., 2001). In this way, distinct populations of neurons can be integrated into transient task-driven assemblies (Buzsáki, 2010) for initiating motor activity (Baker et al., 1999; Parkkonen et al., 2015) or attentional, and memory processes (Jensen et al., 2007; Schroeder & Lakatos, 2009). In summary, coordinated oscillations enable information-coding ensembles to effectively communicate across short and long spatial ranges and with systematic timeframes (Bree et al., 2025). Yet, there are no self-sustained smooth oscillations in the LFP. Instead, signals are composed of transient bursts, and action potentials travel within bursts of LFP oscillations (Bartolo & Merchant, 2015; Cadena-Valencia et al., 2018; Chen et al., 2023). Hence, it is evident that more work is needed to bridge the gap between bursting local oscillations, discharge rate information of single cells, and neural population manifolds as substrates of beat extraction and production. Various large-scale, low-dimensional theoretical models have offered to explain how collective dynamics organize neural ensembles into coherent functional units at different levels of neurophysiological detail (Breakspear, 2017). Neural manifolds can inform such models and provide insight into what neural dynamics would be necessary for human-like self-sustained beat perception and production that unfolds over for indefinite periods of time.

## Acknowledgments

We thank Luis Prado, Raul Paulín, Maria Antonieta Carbajo and Juan Ortíz for their valuable technical assistance The work was supported by UNAM-DGAPA-PAPIIT: IG200424, Consejo Nacional de Humanidades, Ciencia y Tecnología (CONAHCYT) Grant: CBF-2025-G-89 to HM, and NIH P20GM109090 to DD.

